# The heme-scavenger, hemopexin, protects against fungal lung injury by mitigating NETosis: an experimental and computational study

**DOI:** 10.1101/2024.08.06.606866

**Authors:** Ganlin Qu, Henrique A. L. Ribeiro, Angelica L. Solomon, Luis Sordo Vieira, Yana Goddard, Nickolas G. Diodati, Arantxa V Lazarte, Matthew Wheeler, Reinhard Laubenbacher, Borna Mehrad

**Author notes:** Corresponding author: Borna Mehrad Box 100225 University of Florida Gainesville, FL 32610-0225. The authors have declared that no conflicts of interest exist.

## Abstract

Invasive aspergillosis is characterized by lung hemorrhage and release of extracellular heme, which promotes fungal growth. Heme can also mediate tissue injury directly, and both fungal growth and lung injury may induce hemorrhage. To assimilate these interdependent processes, we hypothesized that, during aspergillosis, heme mediates direct lung injury independent of fungal growth, leading to worse infection outcomes, and the scavenger protein, hemopexin, mitigates these effects. Mice with neutropenic aspergillosis were found to have a time-dependent increase in lung extracellular heme and a corresponding hemopexin induction. Hemopexin deficiency resulted in markedly increased lung injury, fungal growth, and lung hemorrhage. Using a computational model of the interactions of *Aspergillus*, heme, and the host, we predicted a critical role for heme-mediated generation of neutrophil-extracellular traps in this infection. We tested this prediction using a fungal strain unable to grow at body temperature, and found that extracellular heme and fungal exposure synergize to induce lung injury by promoting NET release, and disruption of NETs was sufficient to attenuate lung injury and fungal burden. These data implicate heme-mediated NETosis in both lung injury and fungal growth during aspergillosis, resulting in a detrimental positive feedback cycle that can be interrupted by scavenging heme or disrupting NETs.

## Introduction

Invasive pulmonary aspergillosis is a common fungal pneumonia of immunocompromised hosts, and carries a high mortality (1). The spores of the fungus are ubiquitous in air and are inhaled daily by all humans. Normal hosts clear these inhaled spores without becoming ill but, in individuals with impaired immunity, the spores germinate into hyphal filaments that penetrate the lung epithelium, cause pneumonia, and can then disseminate to other organs. Aspergillosis is becoming more common and more resistant to antifungal drugs, prompting the World Health Organization to categorize it as a “critical priority” for research and public health (2).

Pulmonary aspergillosis has long been recognized to result in lung hemorrhage, both in the human infection and in rodent models, thought to result from the physical disruption of blood vessel walls by the invading fungal hyphae (3–5). This hemorrhage causes local hemolysis: the lysis of extravascular red blood cells releases hemoglobin into the extracellular space, which is then degraded to release extracellular heme (6, 7). We recently reported that extracellular heme is utilized by *Aspergillus* as a source of iron, rendering the pathogen more virulent and worsening the outcome of the infection (5). In addition, however, extracellular heme is an endogenous danger signal – an alarmin – that interacts with the host cells via multiple context-dependent mechanisms. Conversely, the host mitigates these injurious effects by scavenging extracellular heme with the liver-derived plasma acute-phase protein, hemopexin (6). The role of extracellular heme and its clearance in progression of aspergillosis and associated tissue injury is only beginning to be understood.

A significant challenge to defining this biology is its complexity: The outcome of aspergillosis is determined by the interplay of multiple processes that unfold over time, including fungal growth, fungal killing by the host, hemorrhage and local hemolysis, heme sequestration, the effects of heme on both the fungus and host cells, and the development of lung injury – which can, itself, cause more hemorrhage. These mechanisms relate to each other by feedback, feedforward, and redundant loops that can progress in parallel or in series – thus producing a complex network. Mechanistic computational modeling is a powerful tool that is well-suited to the systematic integration of such complex biological mechanisms, and is more efficient than serial testing of hypotheses focused on individual nodes in the network. A computational model, based on experimental data, provides a dynamic simulation of the system that can be used to generate novel hypotheses, and serve as an *in silico* laboratory to assess predicted outcomes of hypothetical scenarios. We previously generated a multi-scale agent-based model of aspergillosis, explicitly representing the infected lung in 3 dimensions, within which cells reside and move, and interact with different molecular species (8–10). In the current manuscript, we used an expansion of this computational model, in concert with experimental approaches, to test the hypothesis that, during aspergillosis, heme mediates direct lung injury independent of fungal growth, leading to worse infection outcomes, and the scavenger protein, hemopexin, mitigates these effects.

## Results

### Progressive increase in heme and hemopexin during invasive pulmonary aspergillosis

We began by measuring the lung extracellular heme content during invasive aspergillosis in neutropenic wildtype mice, and found a time-dependent increase in bronchoalveolar lavage (BAL) extracellular heme with aspergillosis (Figure 1A), which paralleled the extent of lung hemorrhage (5). The transcription of hemopexin, a liver-derived acute phase protein that scavenges extracellular heme, was similarly upregulated and correlated with the concentration of hemopexin in plasma (Figure 1B-C). The lung hemopexin concentration also increased over time, but was two orders of magnitude lower than plasma (Figure 1C), consistent with the diffusion of plasma hemopexin into the lungs. To test whether plasma hemopexin scavenges extracellular heme in infected lungs, we measured plasma hemopexin concentration after intra-pulmonary administration of exogenous heme to animals with experimental aspergillosis. Intra-pulmonary administration of heme on day 1 of infection resulted in lower plasma hemopexin on day 2 (Figure 1D). Taken together, these data suggest that, during invasive aspergillosis, hemopexin synthesis is upregulated in the liver, reaches the lungs via the circulation, where it scavenges extracellular heme.

**Figure 1.**
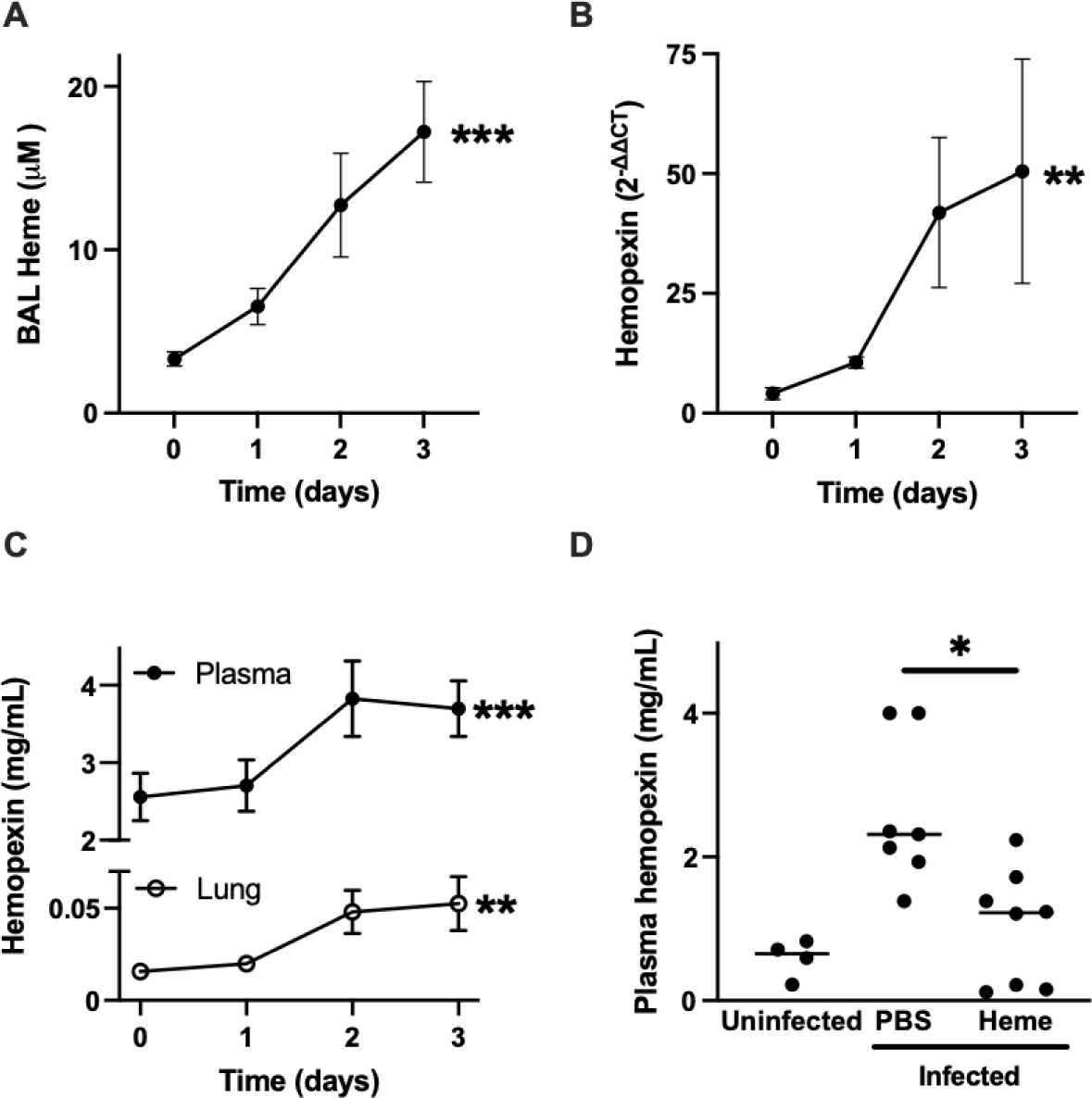
Lung heme content and hemopexin during invasive pulmonary aspergillosis. (A) The concentration of heme in bronchoalveolar lavage (BAL) over the first 3 days of invasive aspergillosis. (B-C) Time-series of liver hemopexin transcription and plasma and Lung hemopexin concentrations. (D) Effect of exogenous administration of heme on plasma hemopexin during aspergillosis. Intra-pulmonary administration of heme on day 1 of infection resulted in lower plasma hemopexin on day 2 of infection. In panels A-C, values represent mean and standard error of the mean (SEM) of n = 8-11 animals per time point or per group in each panel, and time 0 indicates neutropenic but uninfected animals. In panel D, dots represent individual animals and horizontal lines represent medians. *, **, and *** denote p values of <0.05, <0.01, and <0.001, respectively.

### Hemopexin is essential to host defense in invasive aspergillosis

To determine the contribution of hemopexin to host defense in aspergillosis, we next infected neutropenic wildtype and hemopexin-deficient mice with *Aspergillus* conidia. Hemopexin-deficient mice had dramatically increased mortality (Figure 2A) and, on histology, displayed greater extent of leukocyte infiltration into the lungs and more extensive invasion of lung tissue with fungal hyphae than wildtypes (Figure 2B). To quantify the extent of lung injury, we measured BAL albumin and found that hemopexin-deficient animals have normal lung barrier integrity at baseline but increased permeability during the infection (Figure 2C). This increased degree of lung injury in hemopexin-deficient mice was associated with higher lung fungal burden, as measured by the lung content of (1,3)-β-glucan, a major fungal cell wall carbohydrate released from the growing hyphae (11); and lung colony-forming units (CFU) (Figure 2D-E). Furthermore, administration of hemopexin to infected hemopexin-deficient mice resulted in lower lung fungal burden (Supplemental Figure S1).

**Figure 2.**
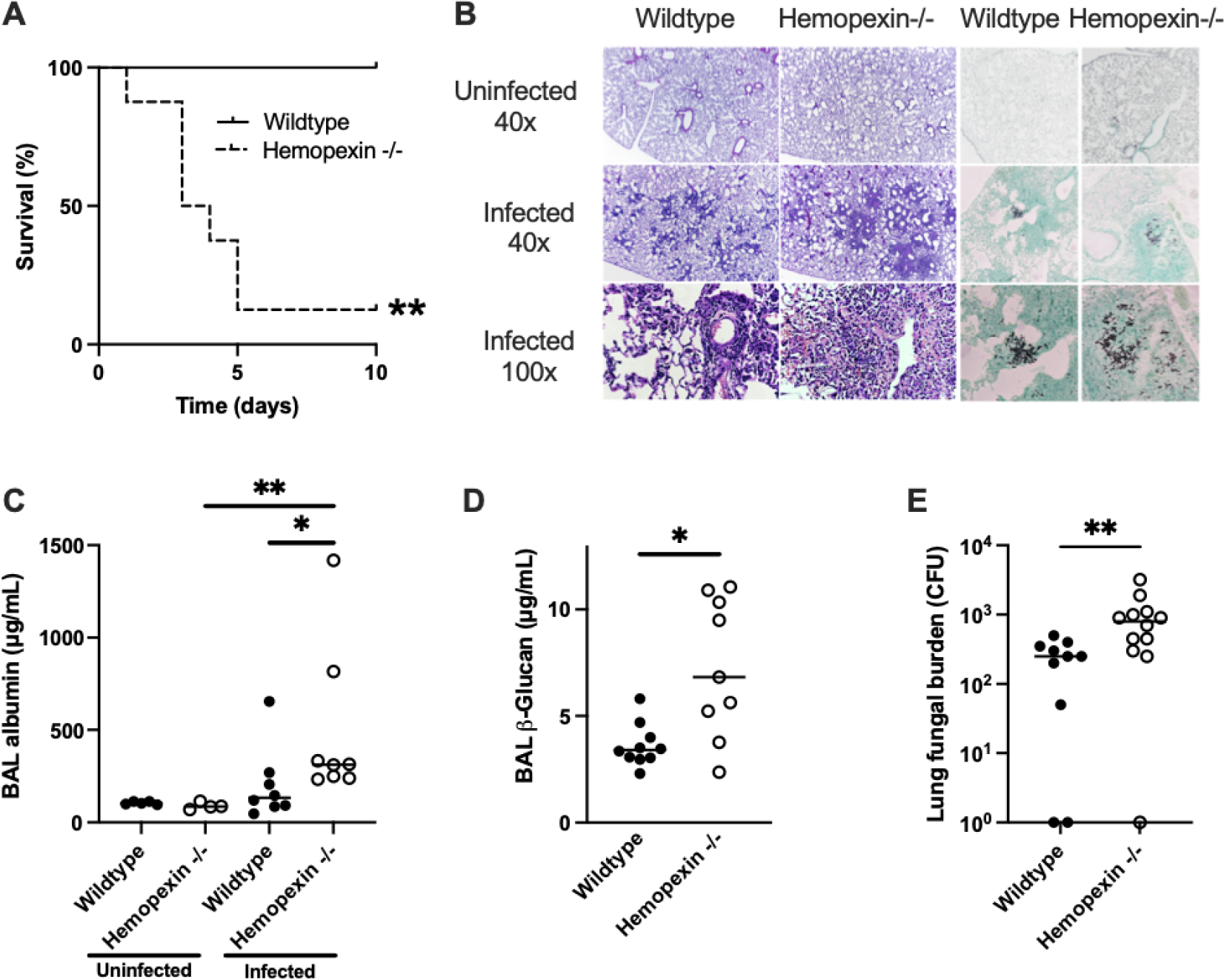
Protective role of hemopexin in invasive pulmonary aspergillosis. (A) Survival of neutropenic wildtype and hemopexin-deficient mice with invasive aspergillosis. n=6-8 per group. (B) Lung histology of wildtype and hemopexin-deficient mice on day 3 of infection or without infection. Hematoxylin and eosin and Gomori methenamine silver stains. (C) Extent of lung injury, measured as BAL fluid albumin concentration, in neutropenic mice with or without aspergillosis. (D-E) Lung fungal content, measured as BAL fluid β-glucan content and lung colony-forming units (CFU) in neutropenic mice with aspergillosis. In panels C-E, dots represent individual animals and horizontal lines represent medians. Since zero cannot be depicted on a log scale, animals with no fungal growth are depicted as having 1 CFU in panel E. * and ** denote *p* values of <0.05 and <0.01, respectively.

We next assessed the effect of hemopexin on local lung hemolysis during aspergillosis. As expected, there was no difference between uninfected wildtype and hemopexin-deficient mice in lung heme and hemoglobin content, and lung heme content was higher in infected hemopexin-deficient mice as compared to wildtypes (Figure 3A). This hemolysis was localized to the lung, since the concentration of blood free heme did not differ between groups (Figure 3B). Surprisingly, the lung concentration of extracellular hemoglobin was also elevated in the lungs of infected hemopexin-deficient animals (Figure 3C), suggesting that the presence of extracellular heme in the infected lung promotes further hemorrhage.

**Figure 3.**
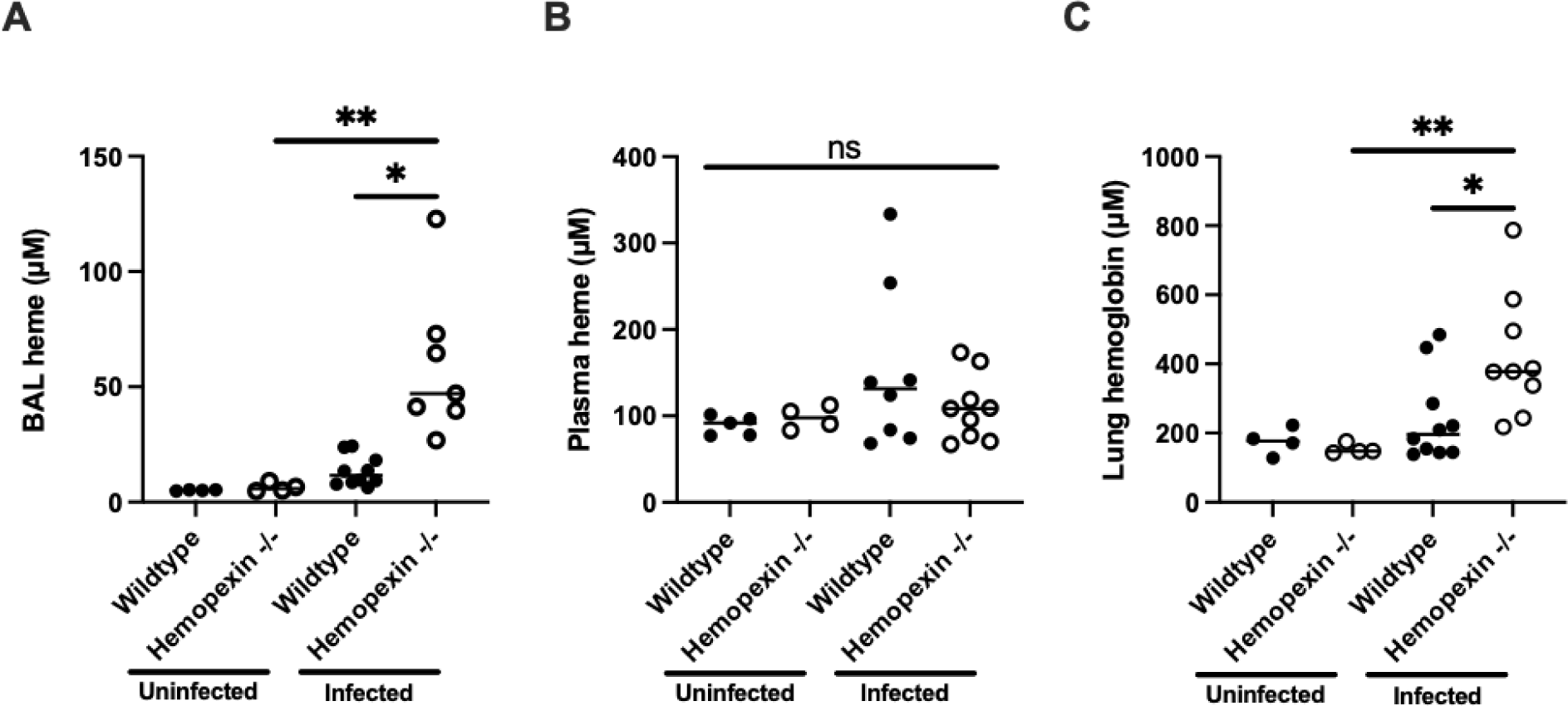
The effect of hemopexin on lung heme and hemoglobin accumulation in aspergillosis. (A-B) Bronchoalveolar lavage and plasma heme concentrations in wildtype and hemopexin-deficient neutropenic mice with or without aspergillosis. (C) Lung hemoglobin concentration in wildtype and hemopexin-deficient neutropenic mice with or without aspergillosis. Dots represent individual animals and horizontal lines represent medians. * and ** denote *p* values of <0.05 and <0.01, respectively; ns, no significant difference.

### Computational modeling of the role of heme in pulmonary aspergillosis

We reasoned that several mechanisms can result in the increased lung hemorrhage in hemopexin-deficient mice during aspergillosis – these mechanisms include the effects of extracellular heme on the fungus, promoting its growth and pathogenicity; the effects of extracellular heme on the host, promoting lung injury; or a combination of these. To capture these possibilities, we incorporated several new mechanisms, including hemorrhage and heme release, into a previously generated computational model of the immune response against *A. fumigatus* (9). The new model (a simplification of which is presented in Figure 4A) was constructed and parameterized using the published literature and data from the current manuscript (see Supplemental Data for details). We incorporated neutrophil extracellular traps (NETs) into the computational model, since neutrophils are pivotal in host defenses against *Aspergillus*, both heme and *Aspergillus* induce NET formation (12, 13), and NETs can mediate lung injury (14–16).

**Figure 4.**
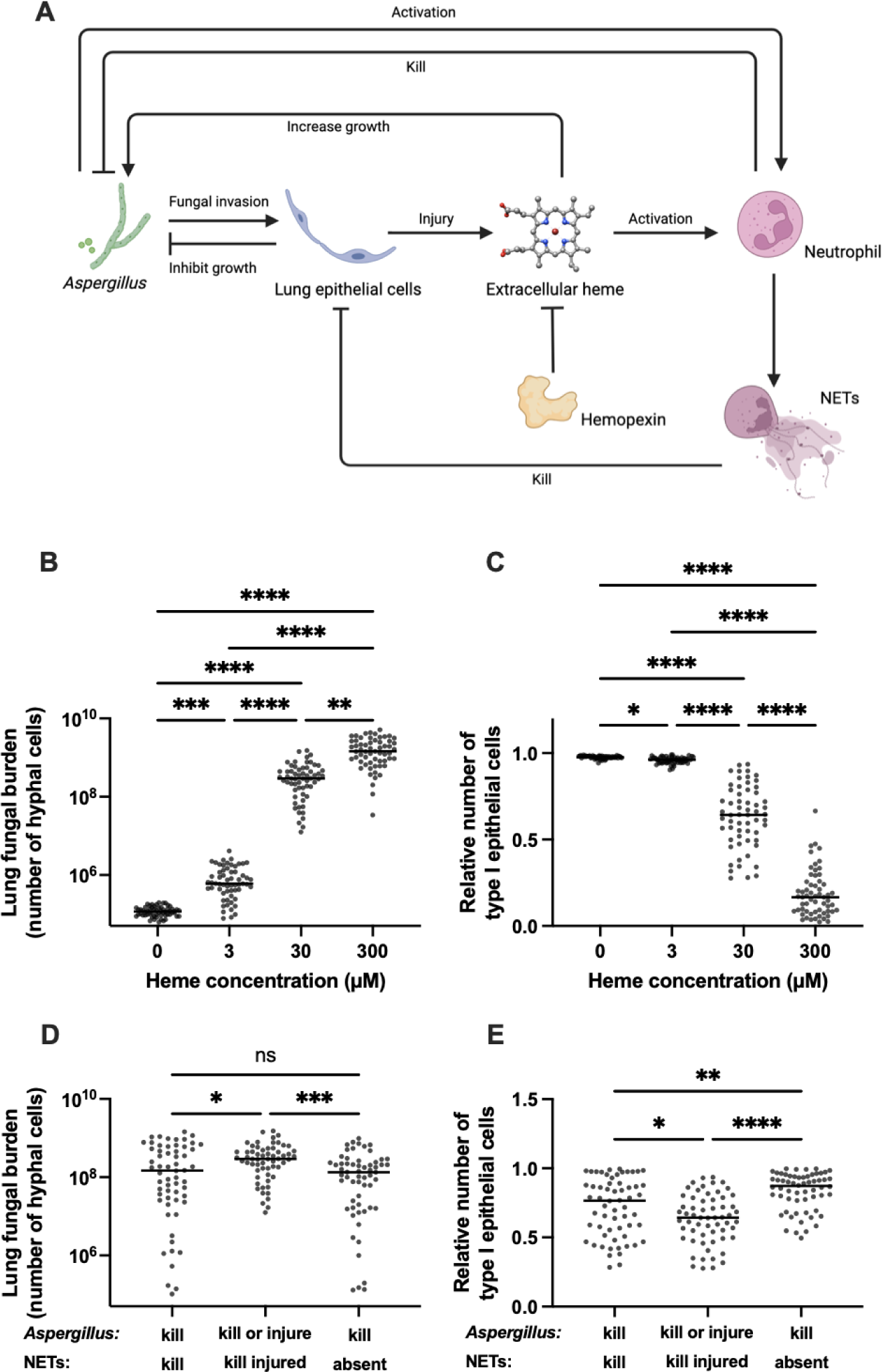
Computational model of the role of heme in aspergillosis. (A) Schematic representation of key mechanisms incorporated into the mathematical model. Image made in Biorender. (B-C) the predicted effect of different concentrations of alveolar extracellular heme on lung fungal load and lung injury. Lung CFU was represented as the number of hyphal fragments between septa, and lung injury is represented as death of type-I alveolar epithelial cells compared to baseline representation of the lung. (D-E) Comparison of the predicted effects of neutrophil extracellular traps on lung fungal burden and injury, in the presence of 30µM heme. X-axis labels describe the model rules regarding effects of *Aspergillus* and NETs on type-I alveolar epithelial cells. Dots represent results of simulation runs and horizontal lines represent medians. *, **, and **** denote *p* values of <0.05, <0.01, and <0.0001 respectively; ns, no significant difference.

Using the computational model, we assessed the effects of lung extracellular heme on the outcome of the infection by running 48 hours of simulated infection in partially neutropenic mice with differing concentrations of extracellular heme in the alveoli. We chose the range of simulated alveolar heme concentrations based on Figures 1A and 3A, accounting for dilution of alveolar contents during bronchoalveolar lavage. The model predicted a positive correlation between alveolar heme concentrations and lung fungal burden (Figure 4B). The model further predicted that increasing concentrations of intra-alveolar heme resulted in progressively greater degree of lung injury, represented in the model by the loss of type I alveolar epithelial cells (Figure 4C).

We next assessed the predicted effect of NETs during the infection, by comparing the predicted effect of 3 hypothetical rules on the loss of type I alveolar epithelial cells (and hence the development of lung injury): In the first condition, NETs and hyphae could each kill epithelial cells independently. In the second condition, hyphae could kill epithelial cells independently, and NETs could only synergistically kill epithelial cells that had previously been injured by hyphae. In the third condition, no NETs were present, and epithelial cell killing was only mediated by hyphae. As expected, the model predicted that synergistic killing results in the most, and absence of NETs resulted in the least, alveolar epithelial cell death. Interestingly, the model also predicted that synergistic killing will result in higher lung fungal burden that was mitigated if NETs were not present (Figure 4D-E).

### Hemopexin protects against aspergillosis-induced lung injury independent of fungal growth

The mathematical model predicted that extravascular heme in the alveolus mediates lung injury both by promoting fungal growth and, independently, via a direct effect on the host. To separate the effect of extracellular heme on the fungus from its predicted effects on the host, we next used the Δ*CgrA* temperature-intolerant mutant strain of *Aspergillus*, which grows at room temperature but not at 37°C, and is hence avirulent *in vivo* (17). We cultured the Δ*CgrA* to the germling stage at room temperature, and then used these in intra-pulmonary challenges in neutropenic mice. As expected, neither wildtype nor hemopexin-deficient neutropenic mice developed appreciable lung injury after challenge with Δ*CgrA* germlings (Figure 5A). In contrast, when Δ*CgrA* germlings were co-administered with intrapulmonary heme, hemopexin-deficient animals developed lung injury in response to the challenge (Figure 5A). To assess whether the latter effect is mediated merely by the intra-pulmonary administration of exogenous heme to hemopexin-deficient animals, we challenged hemopexin-deficient animals with Δ*CgrA* germlings, heme alone, or both, and found that the combination of heme and Δ*CgrA* germlings was necessary to the development of lung injury in these hosts (Figure 5B). Finally, we found similar results when we challenged hemopexin-deficient mice with killed wildtype *Aspergillus* germlings (Figure 5C). Taken together, these data indicate that, independent of fungal growth, intra-alveolar extracellular heme during aspergillosis mediates lung injury, and hemopexin provides protection against this injury – consistent with the prediction of the computational model.

**Figure 5.**
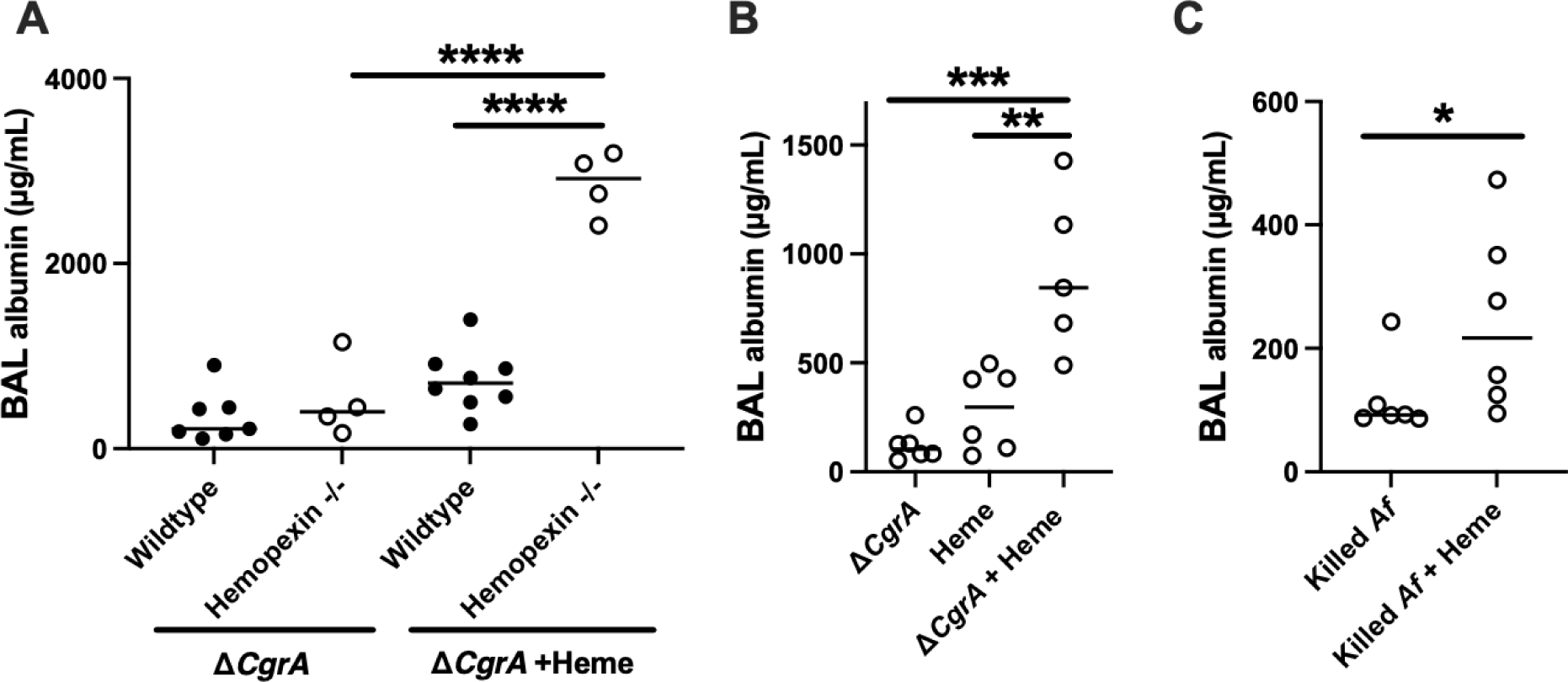
The role of heme in the induction of lung injury in neutropenic aspergillosis. (A) Lung injury in neutropenic wildtype and hemopexin-deficient mice challenged with germlings of Δ*CgrA* (temperature intolerant) strain, with or without intrapulmonary administration of heme. (B) Effect of exogenous heme on lung injury in neutropenic hemopexin-deficient mice challenged with Δ*CgrA* germlings, heme, or both. (C) Effect of exogenous heme on lung injury in neutropenic hemopexin-deficient mice challenged with ethanol killed *Aspergillus* germlings, with or without heme. Dots represent individual animals and horizontal lines represent medians. *, **, ***, and **** denote *p* values of <0.05, <0.01, <0.001, and <0.0001 respectively.

### Heme mediates poor outcomes in aspergillosis by inducing NETosis

We next sought to determine the mechanism by which heme mediates lung injury independently of fungal growth. While heme, acting as an alarmin, can directly affect the host in several ways, the mathematical model predicted a detrimental role in heme-mediated NETosis during aspergillosis (Figure 4D-E). We began to test this prediction by measuring the number of neutrophils and the quantity of NETs in wildtype mice with aspergillosis. We found that, in this partially neutropenic animal model, the number of alveolar neutrophils peaked on day 1 and decreased thereafter; in contrast, the quantity of NETs peaked on day 1 but plateaued thereafter (Figure 6A) – consistent with the notion that neutrophils arriving in the lung on days 2-3 of the infection rapidly undergo NETosis.

**Figure 6.**
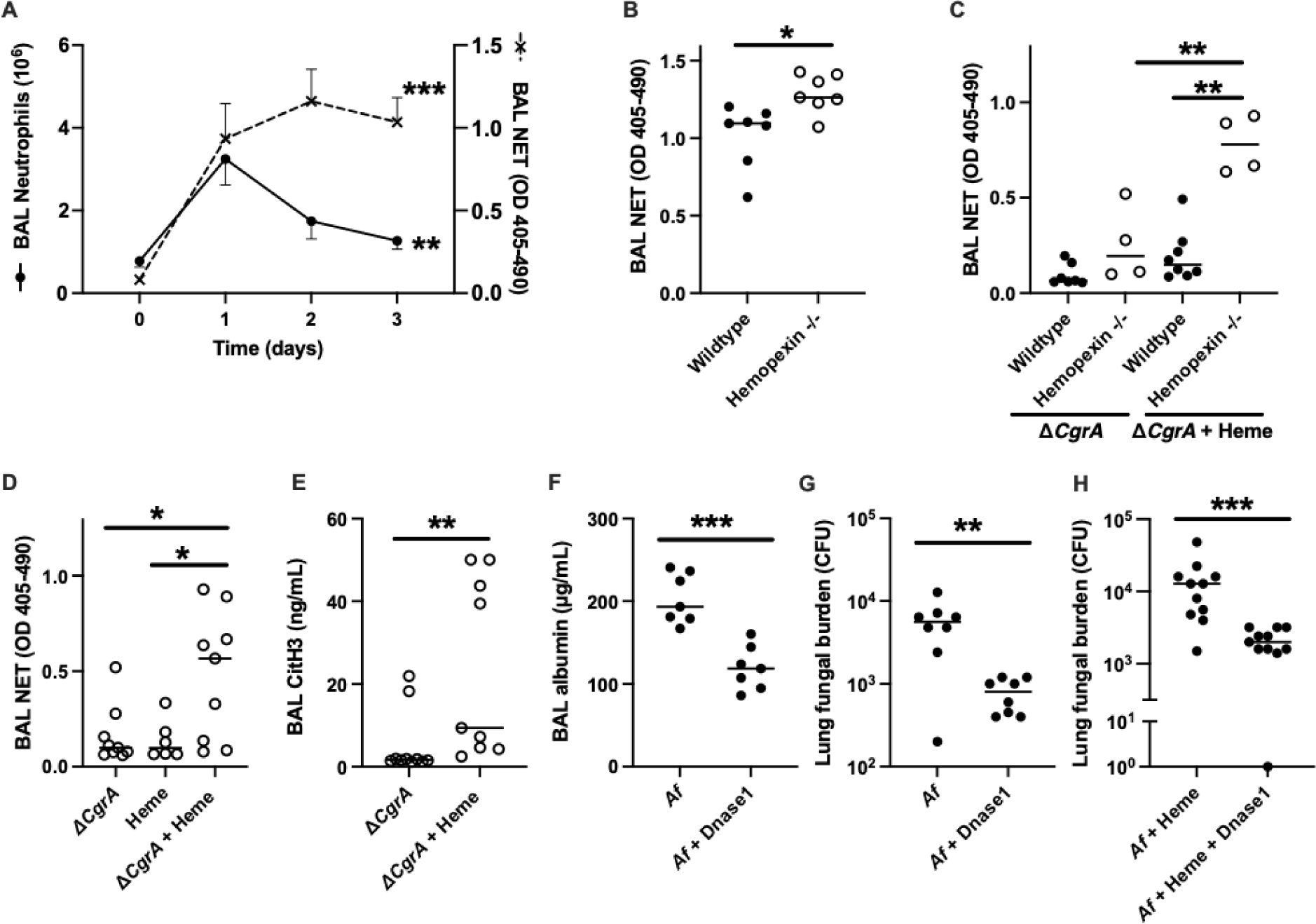
NET formation and resulting lung injury induced by heme which is attenuated by hemopexin. (A) Number of neutrophils and level of NET in the lung of wild type neutropenic mice during invasive pulmonary aspergillosis. (B) BAL NET level of neutropenic mice on day 3 post *Aspergillus* infection. (C) NET level in the plasma and BAL of neutropenic mice after challenge with Δ*CgrA* germlings with or without exogenous heme. (D-E) Effect of exogenous heme on lung NET formation in neutropenic hemopexin-deficient mice challenged with Δ*CgrA* germlings. (F-H) The effect of DNAse treatment on lung injury and fungal burden in wildtype mice with invasive aspergillosis. In F-G, DNAse-1 was administered on day 2 and measurements were taken on day 3 of infection. In H, Dnase1 was administered after 12h and measurements were taken 24h after onset of infection. Since zero cannot be depicted on a log scale, a mouse with no fungal growth is depicted as having 1 CFU in panel H. In panel A, values represent mean ± SEM of n = 5-8 animals per group per time point, and time 0 refers to uninfected neutropenic animals. In panels B-H, dots represent individual animals and horizontal lines represent medians. *, **, and *** denote *p* values of <0.05, <0.01, and <0.001, respectively.

To determine the contribution of heme and hemopexin to NET production during aspergillosis, we next compared BAL NET quantity in wildtype and hemopexin-deficient mice with invasive aspergillosis, and found higher NET levels in hemopexin-deficient mice (Figure 6B). To determine if this observation is attributable to higher fungal burden in hemopexin-deficient animals (Figure 2D-E) or the direct effect of heme on the host, we next compared BAL NET levels in wildtype and hemopexin-deficient mice, challenged with Δ*CgrA* germlings, alone or with intrapulmonary heme. Similar to the degree of lung injury under these conditions (Figure 5A), hemopexin-deficient mice generated more intra-alveolar NETs (Figure 6C), an effect that required challenge with both Δ*CgrA* germlings and heme (Figure 6D). Analogously, intrapulmonary administration of exogenous heme to wildtype mice infected with wildtype *Aspergillus* also resulted in greater lung NET content (Supplemental Figure S2).

A key feature of NET formation is the citrullination of histones contained in the expelled chromatin, a process mediated by peptidylarginine deiminase enzymes (18). To assess this feature of NETosis in aspergillosis, we compared the concentration of citrullinated histone-3 in the alveoli of hemopexin-deficient mice challenged with Δ*CgrA* germlings, with or without heme. In agreement with the NET assay, we found higher concentrations of citrullinated histone-3 in the lungs of animals challenged with both Δ*CgrA* germlings and heme (Figure 6E). Peptidylarginine deiminase-4 (PAD4, one of five peptidylarginine deiminase enzymes) is highly expressed in neutrophils, localizes to the nucleus, and is necessary to NET formation in some disease contexts (19, 20), but not others (21, 22). To determine whether this mechanism is operational in heme-mediated NETosis in aspergillosis, we compared the extent of lung injury between wildtype and PAD4-deficient mice in the context of neutropenic invasive aspergillosis. We found no difference in BAL albumin concentration or quantity of NETs between these groups (Supplemental Figure S3), indicating that PAD4 is dispensable to NETosis in the context of aspergillosis.

The mathematical model predicted that the absence of NETs during aspergillosis will result in less lung injury and, unexpectedly, lower fungal burden (Figure 4D-E). To test these predictions, we examined the effect of intra-pulmonary administration of DNAse-1 to wildtype mice during invasive aspergillosis. As expected, this treatment resulted in lower concentration of NETs in the BAL (Supplemental Figure S4). Consistent with the predictions of the mathematical model, DNAse treatment also resulted in attenuated lung injury and lower lung fungal burden in wildtype mice (Figures 6F-G). Since our earlier data showed that lung extracellular heme was critical to NETosis during aspergillosis (Figures 6C-E), we tested the effect of DNAse-1 treatment in invasive aspergillosis in the context of exogenous administration of heme, and again found a lower lung fungal burden in treated animals (Figure 6H). Collectively, these data implicate heme-mediated NETosis in both lung injury and fungal growth during pulmonary aspergillosis.

## Discussion

The association of aspergillosis with hemorrhage has been documented since the initial descriptions of the infection in the 19th century (23). Histopathologically, *Aspergillus* hyphae invade and disrupt blood vessels, causing intra-alveolar hemorrhage and hemorrhagic and coagulative necrosis in autopsy series (24). We previously reported that *Aspergillus* acquires heme from the abundant extracellular hemoglobin made available by tissue hemorrhage and lysis of red blood cells, rendering it more virulent (5), consistent with the critical dependence of *Aspergillus* species on iron acquisition for their growth (25, 26). In the current work, we found that extracellular heme, independent of its effect on the fungus, also mediates NET formation, which both induces lung injury and impairs fungal killing, and that these effects are mitigated by the heme scavenger hemopexin.

Erythrocytes are the most abundant cell type in the body (27), and their premature destruction – hemolysis – occurs in diverse pathological settings. Following release from erythrocytes, the tetrameric ɑ_2_β_2_ hemoglobin molecules dissociate to ɑβ heterodimers, which are then bound by the liver-derived acute phase protein, haptoglobin, and scavenged by mononuclear phagocytes via the receptor, CD163 (28). Once the available haptoglobin is depleted, dimeric extracellular hemoglobin is oxidized to methemoglobin and releases its heme moiety. The resulting extracellular heme is bound by hemopexin, and scavenged via CD91, a receptor expressed by multiple cell types (28). Independent of its cause, hemolysis results in inflammation, mediated by the release of damage-associated molecular patterns, among which labile extracellular heme is the best characterized. In addition, both dimeric hemoglobin and heme induce oxidative stress (29). These mechanisms have been investigated thoroughly in the context of classical hemolytic conditions, such as hemoglobinopathies (30), but may also be relevant to a broad array of other diseases, given the abundance of erythrocytes and prevalence of hemolysis in different pathologic states.

The above mechanisms have been implicated in lung injury caused by systemic hemolysis, for example in sickle cell acute chest syndrome and non-antibody-mediated transfusion-related lung injury (30, 31). Systemic hemolysis occurs in numerous infections by several mechanisms, namely direct pathogen invasion of erythrocytes, elaboration of hemolysins, and immune-mediated processes (32). Many microbes, including *Aspergillus*, can also take up and use heme as a nutrient (33, 34), but the mechanistic contribution of hemolysis to microbial pathogenesis is defined in only a few infections, most notably in malaria and sepsis (35, 36). The current work adds to this literature by defining the role of local (but not systemic) hemolysis in aspergillosis, via a positive feedback loop in which lung hemorrhage and release of extracellular heme both promotes the growth of the pathogen and mediates further lung injury, thus causing further hemorrhage and local hemolysis.

Formation of NETs is a canonical effector function that is key to both neutrophil antimicrobial effects and inflammatory tissue damage (37). Neutrophils release NETs in response to many pathogens, including *Aspergillus* (12, 38). While NET formation is antimicrobial against many pathogens, the role of neutrophil NETs during aspergillosis remains undefined: Specifically, NETs do not improve neutrophil killing of *Aspergillus* (39–41), and reports of the effect of NETosis on inhibiting fungal growth are conflicting (41, 42). On the other hand, NET formation in the lungs can damage the alveolar epithelium and endothelium, and propagates lung injury (43, 44). Finally, extracellular heme, acting as an alarmin, is a potent inducer of neutrophil NETs *in vitro* and in several disease models (13, 45–47). Our data add to this literature by implicating extracellular heme as a mediator of NETosis during aspergillosis, and identifying NETosis as maladaptive and harmful to the host during this infection. In addition to mitigating lung injury, disruption of NETs in our model also resulted in lower lung fungal burden. As suggested by the computational model, we speculate that lung injury may enhance lung hemorrhage and heme release, thus promoting fungal growth. Furthermore, injury to alveolar epithelial cells and disruption of their antimicrobial functions may further impair fungal clearance.

Computational modeling can help in understanding complex biological systems, by incorporating multiple processes and intertwined feedback loops, and serve as a virtual laboratory to explore possible mechanisms and generate hypotheses to test experimentally. In the current work, we expanded a previously published model that includes many of the components involved in the early immune response to *A. fumigatus* (9), incorporating additional mechanisms to examine the role of heme in this infection. From a modeling perspective, this work advances the field in two ways: first, it demonstrates how a mathematical model of a disease can be used effectively for the discovery of new biology and the efficient testing of experimental hypotheses *in silico* before taking them into the laboratory. Second, it provides an example of how the modular design of the model simplifies its expansion, by allowing the addition of new mechanisms (48). We anticipate that, over time, the model will include many aspects of the immune response to a respiratory infection and for any given question, a sensitivity analysis of model parameters can identify model features that are dispensable in the particular context of the question (49) – thus allowing the model to be adapted to many different investigations.

We conclude that labile extracellular heme released from local hemolysis during aspergillosis is at the nexus of a series of pathogenic mechanisms, including enhancing fungal growth and, separately, induction of lung injury via NETosis, with hemopexin acting as a potent defense against these processes. These findings suggest possible avenues for future research. These include mechanistic studies of other erythrocyte-derived alarmins, and examining the effects of heme on cells other than neutrophils during aspergillosis. Finally, host-centered interventions, such as administration of heme- or hemoglobin-scavengers and inhibition or disruption of NETs, may augment existing therapies for aspergillosis.

## Materials and methods

### Sex as a biological variable

Since male and female animals have similar outcomes in invasive aspergillosis, this study was performed in both male and female animals, and aggregate findings are reported.

### Preparation and administration of A. fumigatus

*A. fumigatus* strain 13073 (American Type Culture Collection, Manassas, Virginia, USA) was cultured on Sabouraud’s dextrose agar plates at 37°C for 2-3 weeks prior to conidia harvested in 0.1% Tween 80 in phosphate-buffered saline. In some experiments, a temperature-sensitive mutant strain of *A. fumigatus* (ΔCgrA, a generous gift from Dr. David Askew, University of Cincinnati) was cultured at room temperature under phleomycin selection for 6-8 weeks prior to conidial harvest, as described (17). The resulting conidia were filtered through sterile gauze, enumerated using a hemocytometer, and cultured in RPMI-1640 containing 1% Penicillin-Streptomycin (plus 10% FBS for the ΔCgrA strain) in n a shaking incubator until the development of germlings. In some experiments, the resulting germlings were killed by resuspending them in 70% ethanol, as described (50).

### Animals, in vivo procedures, and sample collections

Wild-type, hemopexin-deficient, and PAD4-deficient mice, all on C57BL/6J genetic background, were purchased from Jackson Laboratories (Bar Harbor, Maine, USA), and propagated and maintained in a specific-pathogen free vivarium. Age- and sex-matched 7-12-week-old were used in experiments. Aspergillosis was induced as previously described (5, 50). Briefly, mice were rendered neutropenic by intraperitoneal injection of 400µg of anti-Ly6G mAb (clone 1A8, BioXcell, Lebanon, New Hampshire, USA) 1 day before intratracheal administration of 30µL sterile PBS containing 7.5 x 10^6^ resting conidia or 6 x 10^5^ germlings. In some experiments, hemin (Sigma, St. Louis, Missouri, USA) at 1.5 mg/mL in PBS was administered to mice together with *Aspergillus* via the intra-tracheal route, and again 35µL via the intranasal route on day 1 of infection. In other experiments, 4000 units of DNAse-1 (Roche, Mannheim, Germany) in 35µl of PBS was delivered to each mouse intranasally twice a day for 1 day or 2 days of the infection. Sample collections and histology were performed as previously described (51, 52). In some experiments, 125µg of human hemopexin (Athens Research, Athens, Georgia, USA) in 30µL normal saline was administered intranasally on days 1 and day 2 of infection.

### ELISA, quantitative reverse transcriptase PCR, fungal burden measurements, and flow cytometry

Commercial enzyme-linked immunosorbent assays were used to measure the concentrations of albumin (Crystal Chem, Elk Grove Village, Illinois, USA), hemopexin (Abcam, Cambridge, United Kingdom), and 1,3 β-glucan (Associates of Cape Cod, East Falmouth, Massachusetts, USA), according to the manufacturers’ instructions. Heme Assay Kit (Sigma) was used for the quantification of heme, ELISA kit were used to measure hemoglobin ((Abcam, Cambridge, Massachusetts, USA) and citrullinated histone-3 (Cayman Chemical, Ann Arbor, Michigan, USA) per manufacturers’ instructions. Neutrophil extracellular traps were quantified by measuring neutrophil elastase-DNA complex using an in-house ELISA, as previously described (19), using anti-mouse neutrophil elastase antibody (clone G-2, Santa Cruz Biotechnology, Santa Cruz, California, USA) as capture antibody, and peroxidase-conjugated anti-DNA antibody (clone MCA-33, Roche Applied Science, Penzberg, Germany) as detection antibody.

For quantitative reverse-transcriptase polymerase chain reaction, liver RNA was isolated (RNeasy Plus Mini Kit, Quiagen, Venlo, Netherlands), and complementary DNA synthesized using a reverse-transcription kit (QUANTAbio, Beverly, Massachusetts, USA). Hemopexin transcript was quantified in duplicate using SYBR green (Bio-Rad, Hercules, California, USA). Expression was calculated using the ΔΔC(t) method normalized to peptidylprolyl isomerase A (PPIA), and then relative to expression of uninfected animals, using commercial primers (qMmuCED0041303 and qMmuCID0016484, Bio-Rad, Hercules, California, USA). Reactions were performed on a iQ5 Thermal Cycler (Bio-rad) using the following settings: 2 minutes at 95°C for activation, 5 seconds at 95°C and 30 seconds at 60°C for 40 cycles.

Fungal colony-forming units (CFU) were determined by homogenizing freshly harvested lung tissues in 1 mL distilled water at 50 Hz for 10 minutes (TissueLyser LT, Quiagen), followed by serial 2-fold dilutions of the samples in water and culture, in duplicate, on Sabourad’s dextrose agar plates containing 0.05% Triton X-100. After ∼18h of incubation at 37°C, plates were photographed and CFU enumerated.

Flow cytometry was performed as described previously (51, 53). Briefly, cell suspensions from freshly harvested mouse lung tissue were generated by incubating mice tissues in RPMI-1640 containing 10% FBS, liberase (Sigma), and Dnase (Sigma) for 1 hour at 37°C. After complete digestion, single-cell suspension was collected and stained with neutrophils markers (all from BD Biosciences, San Jose, California, USA or BioLegend, San Diego, California, USA). After live dead staining by Zombie Aqua™ dye and Fc block (anti-CD16/CD32), we characterized neutrophils as anti-CD45–PerCP (clone 30-F11), anti-CD11b–FITC (Clone M1/70), and anti-Ly6G-Brilliant Violet 605 (clone 1A8). Data was collected and analyzed using FlowJo software v. 9.0 (Tree Star, Ashland, Oregon, USA). Neutrophils were identified as CD45+, CD11b+, and Ly6G+ cells (Supplemental Figure S5).

### Computational modeling

We expanded a previously published computational model of invasive aspergillosis (9), as described in detail in the Supplemental Data. Briefly, the model uses a three-dimensional toroidal homogeneous space with a volume of 6.4 x 10^-2^ mm^3^, representing a portion of the murine lung. This space is composed of 10 x 10 x 10 voxels, with each voxel representing a 40 x 40 x 40µm mouse alveolus, which at baseline contains type I and type II alveolar epithelial cells and alveolar macrophages. The infection in this space is initialized with 1920 resting *Aspergillus* conidia, which scales to 10^7^ intrapulmonary conidia per mouse, the approximate inoculum size we use experimentally. After 4h of simulated time, the resting conidia begin to swell, and then germinate into hyphae that invade the lung, resulting in recruitment of macrophages and neutrophils. When in contact with hyphae, type I alveolar epithelial cells can be either injured or killed. Type I alveolar epithelial cells can also be killed when in contact with NETs: NETs can kill any healthy cell with a low probability (< 5%), and kill type I alveolar epithelial cells previously injured by hyphae with 100% probability. Death of type I alveolar epithelial cells results in filling of the alveolus with blood and release of extracellular heme. Heme is taken up by hyphae, increasing their growth. Heme contributes to hyphal growth both by providing a source of iron, and the protoporphyrin ring, which independently contributes to fungal growth (5). The type I epithelial cells inhibit fungal growth in a contact-dependent manner. Recruited neutrophils can be activated by fungal (1,3)-β-D-glucan and by heme. Activated neutrophils kill *Aspergillus*, and have a half-life of 6h, after which they undergo NETosis or apoptosis. The resulting NETs are degraded with a half-life of 3h (See Table S1).

### Statistical analysis

All experiments were performed at least twice, and combined data are presented. Data were analyzed using the Prism software (version 10.2; GraphPad, Boston, Massachusetts, USA). Differences between two groups at one time point were analyzed with an unpaired two-tailed Mann-Whitney (nonparametric) test and comparisons between ≥3 groups were performed using Kruskal-Wallis one-way ANOVA with Dunn’s multiple comparison test. Survival comparisons were performed using the Log-rank test. Statistically significance was assigned to a p value less than 0.05.

### Study approval

The animal studies reported in this manuscript were approved by our institutional review board.

### Data availability

Most of the data in this manuscript is presented as individual data points. In instances where only summary data are presented, individual data points are provided in the Supplemental Data Values file. The computer code for the mathematical model is available at https://github.com/NutritionalLungImmunity/jISSnet/tree/master.

## Author contributions

Conception and design: GQ, RL, BM Data collection: GQ, ALS, LSV, YG, MW

Analysis and interpretation: GQ, HALR, ALS, RL, BM Drafting the manuscript: GQ, HALR, RL, BM

Critical revision: ALS, LSV, RL, BM

Final approval of the version submitted: all authors.

