## Supplemental data for "The heme-scavenger, hemopexin, protects against fungal lung injury by mitigating NETosis: an experimental and computational study"

Ganlin Qu (ORCID ID 0009-0000-0161-5144), Henrique A. L. Ribeiro (ORCID ID 0000-0001-9802-5211), Angelica L. Solomon (ORCID ID 0009-0004-8463-3960), Luis Sordo Vieira (ORCID ID 0000-0002-3608-9356), Yana Goddard (ORCID ID 0009-0003-5448-1823), Matthew Wheeler (ORCID ID 0000-0001-7396-5469), Reinhard Laubenbacher (ORCID ID 0000-0002-9143-9451), Borna Mehrad (ORCID ID 0000-0001-5198-065X) [1]

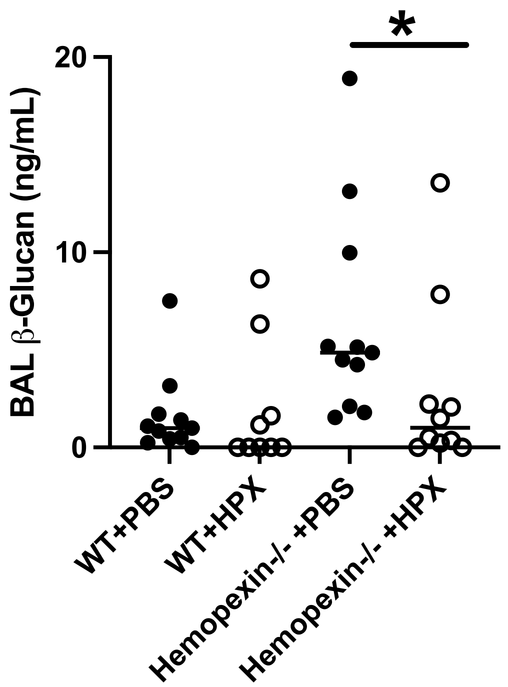


**Figure S1.** The effect of administration of intra-pulmonary hemopexin (HPX), as compared to PBS vehicle, on lung fungal burden (measured as BAL β-glucan concentration) on day 3 of infection. Dots represent individual animals and horizontal lines represent medians. * denotes *p* values of <0.05.

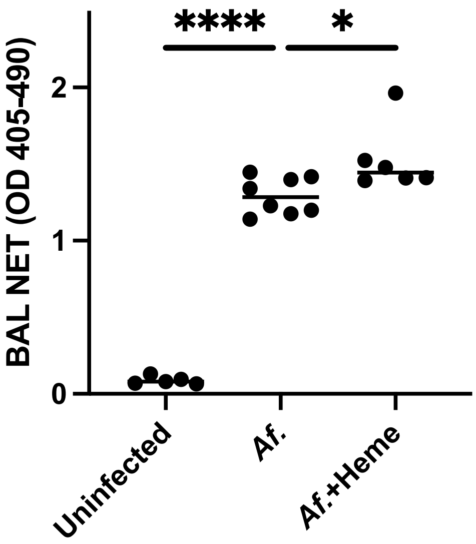


**Figure S2.** The effect of heme on lung NET formation in wildtype mice with neutropenic aspergillosis. BAL NET level of uninfected wildtype mice and *Aspergillus* infected wildtype mice with or without administration of heme. Dots represent individual animals and horizontal lines represent medians. *, and **** denote $p$ values of <0.05 and <0.0001 respectively.

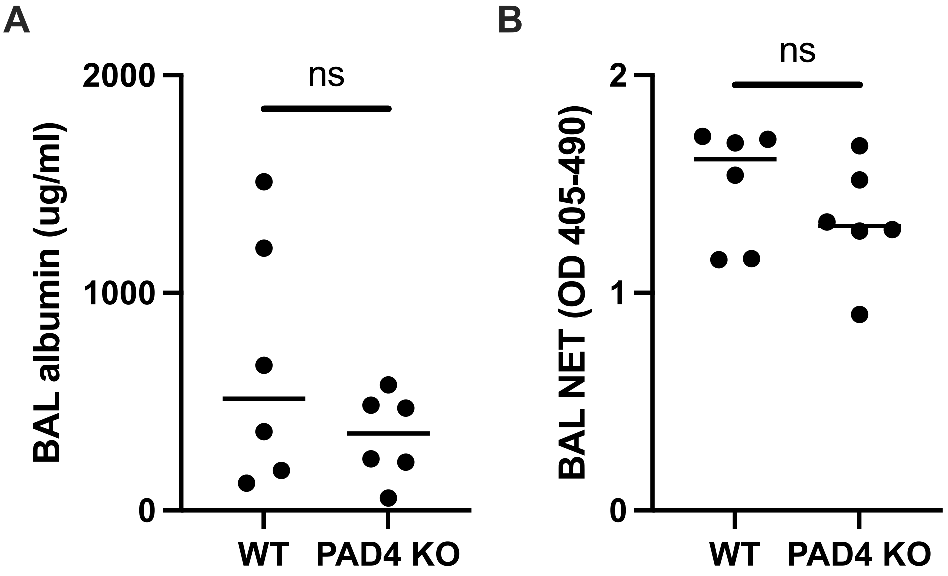


**Figure S3.** Lung injury and NET formation caused by aspergillosis is independent of PAD4. Extent of lung injury, as measured as BAL fluid albumin concentration (panel A) and level of BAL NETs (panel B) in neutropenic mice with aspergillosis. Dots represent individual animals and horizontal lines represent medians. ns, no significant difference.

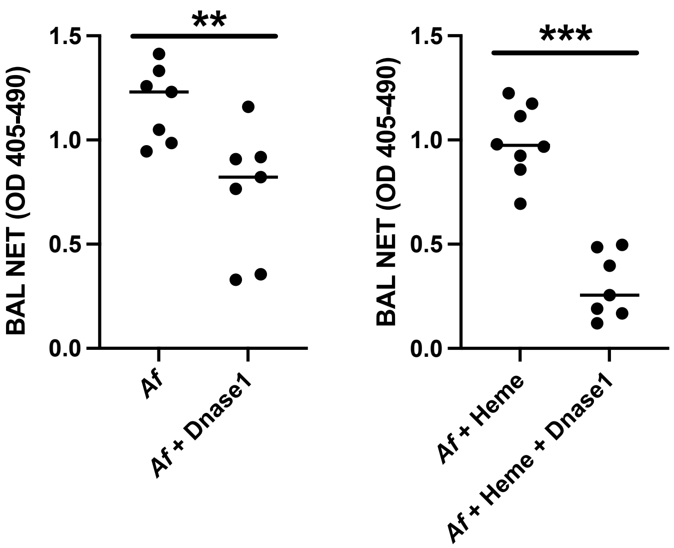


**Figure S4.** Effect of DNAse treatment on lung NET formation in wildtype mice with neutropenic aspergillosis. (A) DNAse-1 was administered on day 2 and BAL NETs measured on day 3 of infection. (B) Dnase1 was administered on day 1 and measurements were taken on day 2 of infection. Dots represent individual animals and horizontal lines represent medians. ** and *** denote *p* values of <0.01 and <0.001 respectively.

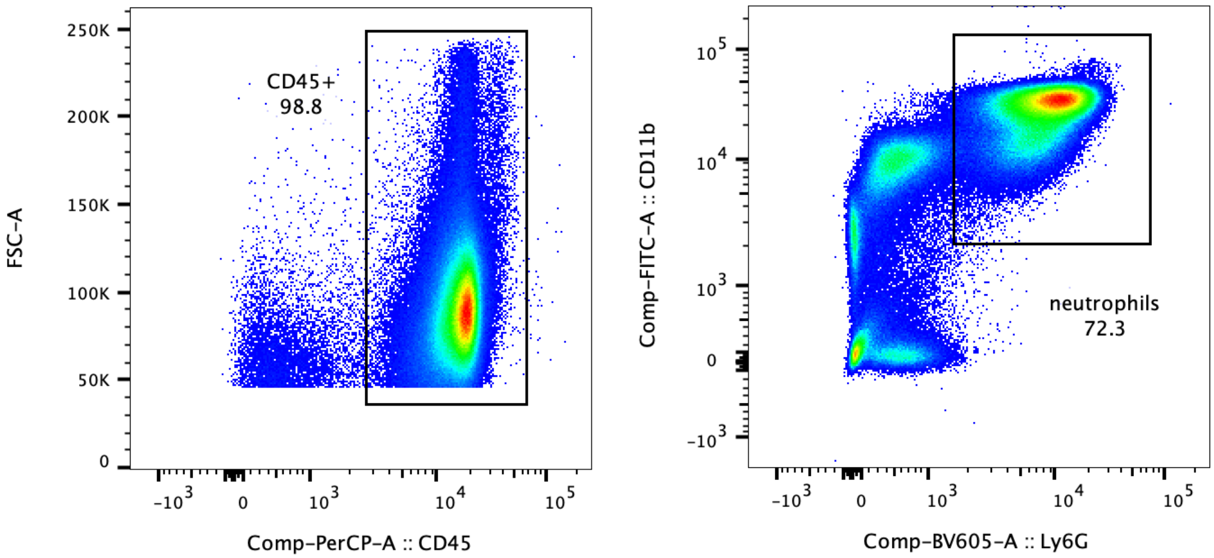


**Figure S5.** Flow cytometry gating strategy of lung neutrophils. Lung cells were first gated on single cells, then live cells, followed by CD45+, CD11b+, and Ly6G+ cells.

### Computational model

Description of the base model

The base model is described in detail in [1]. We provide a general description of this model in this section for completeness: The model contains five cell types: *Aspergillus* (as resting conidia, swollen conidia, or hyphae), neutrophils, macrophages, and types I and II pneumocytes. Macrophages and neutrophils are motile cells and move randomly or biased towards a chemokine gradient, and die with a half-life (Table S1). Contact with hyphae and swollen conidia activates leukocytes and type II pneumocytes, leading them to secrete cytokines (TNF, IL10, CCL4, CXCL2). Leukocytes can phagocytose swollen conidia and kill hyphae. Type I pneumocytes inhibit hyphae elongation by a contact-mediated mechanism. Contact with the fungus and TNF each activate macrophages classically, to an M1 phenotype, whereas IL10, TGF, and apoptotic bodies activate macrophages to an M2 phenotype incapable of killing. Activated neutrophils also degranulate, releasing lactoferrin, a molecule that competes with fungal siderophore, TAFC, for iron.

*Aspergillus* starts as resting conidia. After four hours, it becomes swollen becomes visible for leukocytes and type II pneumocytes, and begins secreting siderophores. Both iron and heme serve as nutrients for fungal growth: with enough iron, *Aspergillus* conidia germinate, and hyphae will elongate and branch. Equation 1 computes the number of iterations for the next 40 um segment of fungi to grow, given the internal concentration of heme and iron. We use the reciprocal of Michaelean kinetics as a phenomenological equation to integrate heme and iron as nutrients that contributing to fungal growth.

$$t = T \cdot \frac{K_I \cdot I + K_H \cdot H + I \cdot H}{H \cdot I} \cdot EPI_{INH}$$

Equation 1: where "t" is the number of iterations until the next 40 um hyphal fragment grows. "T" is the inverse of the growth rate (r). $K_I$ is the iron $K_M$, and $K_H$ is the heme $K_M$. I and H are internal iron and heme concentrations. $EPI_{INH}$ is the growth inhibition by alveolar epithelial cells.

Cytokines and TAFC released into the alveolus diffuse according to a partial differential equation [1], undergo decay (with a half-life), and diffuse into plasma.

Changes from the published base model

Although cells interact with their environment continuously, the experimental data that the model is based on is not continuous. In order for the model to more closely match the available experimental data, we revised the model so that each cell interacts with its environment every 30 minutes instead of every iteration. In every iteration, we select a subset of cells of a type (such as a subset of the macrophage) to interact with other cells and molecules, with any given cell is selected only every 15 iterations (30 minutes). Likewise, the Boolean networks are updated every thirty minutes.

We rewrote the state model from epithelial cells and neutrophils as Boolean networks. Neutrophils are activated by heme or *Aspergillus*. Activated neutrophils release lactoferrin and then change to either apoptotic or NETotic states, whereas non-activated neutrophils only die by apoptosis. Neutrophil apoptosis or NETosis occur with a half-life of 6h. NETs and hyphae kill type I pneumocytes, result in release of extracellular heme into the alveolus. In addition to neutrophils, heme also activates M1 macrophages (described below). Once a type I epithelial cell dies, it results in the appearance of extracellular heme in the alveolus, simulating hemorrhage. The heme quantity is finite and is not replenished and does not diffuse, simulating clotting after hemorrhage. Once the type I pneumocytes die, their inhibition over hyphae elongation is lifted.

We refined the macrophage state model to a Boolean network model based on [104]. We ran the network until it reached equilibrium, and then assessed the macrophage phenotypes. Macrophages with activation of NFKB, STAT1, or STAT5 are classified as M1; STAT6 as M2A; ERK as M2B; and STAT3 as M2C.

Since the quantity of cytokines secreted by neutrophils was minimal in the previous model, we eliminated neutrophil cytokine secretion in this version. We also removed the IL6-hepcidin axis from the model, because this axis had little effect on the model outcome, and experimental evidence from our group showed that hepcidin did not affect infection evolution [69].

**Table S1:** Parameters of the revised mathematical model.

| ID | Parameter | Description | *Value* | Reference |
|---|---|---|---|---|
| 1 | CCL4_QTTY | CCL4 secretion rate by macrophages and type II pneumocytes | $1.79 \times 10^{-20}\ \mathrm{mol \cdot cell^{-1} \cdot h^{-1}}$ | [2,3,4,5,6,7,8,9,10,11,12,13,14,15,16,17,18,19,20,21,22,23,24,25,26,27,28,29,30,31] |
| 2 | CXCL2_QTTY | CXCL2 secretion rate by macrophages and type II pneumocytes | $1.11 \times 10^{-19}\ \mathrm{mol \cdot cell^{-1} \cdot h^{-1}}$ | |
| 3 | TNF_QTTY | TNF secretion rate by macrophage and type II pneumocytes | $3.22 \times 10^{-20}\ \mathrm{mol \cdot cell^{-1} \cdot h^{-1}}$ | |
| 4 | IL10_QTTY | IL10 secretion rate by macrophages | $6.97 \times 10^{-22}\ \mathrm{mol \cdot cell^{-1} \cdot h^{-1}}$ | |
| 5 | TGF_QTTY | TGF secretion rate by macrophages | $1.01 \times 10^{-21}\ \mathrm{mol \cdot cell^{-1} \cdot h^{-1}}$ | |
| 6 | Lac_QTTY | Amount of lactoferrin | $5.36 \times 10^{-18}\ \mathrm{mol \cdot cell^{-1}}$ | [32] |
| 7 | Kd_CCL4 | Kd of the CCL4 receptor | 180 pM | [33] |
| 8 | Kd_CXCL2 | Kd of the CXCL2 receptor | 91.667 pM | [34,35] |
| 9 | Kd_IL10 | Kd of the IL10 receptor | 140 pM | [36,37,38,39] |
| 10 | Kd_TNF | Kd of the TNF receptor | 326 pM | [40,41,42,43,44,45,46,47] |
| 11 | Kd_TGF | Kd of the TGF receptor | 26.5 pM | [48,49,50] |
| 12 | D | Diffusion rate | $850\ \mu\mathrm{m}^2/\mathrm{min}$ | [51,52] |
| 13 | $\lambda$ | Cytokine half-life | 1h | [53,54,55,56,57, 58,59] |
| 14 | TF_CONC | Transferrin concentration | 32.25 μM | [60,61,62] |
| 15 | APO_TF_REL_CON | Apo-transferrin relative concentration | 40% | [62, 63] |
| 16 | TFFE_REL_CON | Mono-ferric transferrin relative concentration | 16.57% | |
| 17 | TFFE2_REL_CON | Di-ferric transferrin relative concentration | 43.43% | |
| 18 | MA_IRON_EXP | Macrophage iron export rate | $1367.30\ \mathrm{M^{-1} \cdot h^{-1}}$ | [64] |
| 19 | MA_IRON_IMP | Macrophage iron import rate | $5.33 \times 10^{-12}\ \mathrm{L \cdot cell^{-1} \cdot h^{-1}}$ | [64] |
| 20 | MA_INT_IRON | Macrophage initial internal iron concentration | $1.0086 \times 10^{-14}\ \mathrm{mol}$ | [62] |
| 21 | TAFC_QTTY | TAFC secretion rate | $1.0 \times 10^{-15}\ \mathrm{mol \cdot cell^{-1} \cdot h^{-1}}$ | [65] |
| 22 | TAFCBI_UPTAKE | uptake rate of TAFC bound to iron | $1.0 \times 10^{-12}\ \mathrm{L \cdot cell^{-1} \cdot h^{-1}}$ | [66,67] |
| 23 | $K_I$ | Michaelian constant for the iron substrate in Equation 1 | 47.43 μM | [68] |
| 24 | $K_H$ | Michaelian constant for the heme substrate in Equation 1 | 979.01 nM | [69] |
| 25 | Kcat_Km_TAFC | Kcat/Km TAFC-Tf iron chelation reaction | $397.77\ \mathrm{M^{-1} \cdot s^{-1}}$ | [63,65] |
| 26 | Kcat_Km_LAC | Kcat/Km lactoferrin-Tf iron chelation reaction | $399.20\ \mathrm{M^{-1} \cdot s^{-1}}$ | [32] |
| 27 | r | Hyphae elongation rate | 80 μm/h | [70] |
| 28 | PR_BR | Hyphae branch probability | 33.3% | [70] |
| 29 | PR_SW | Conidia swelling probability | 0.39% | [71,72] |
| 30 | T_SWELL | Time until conidia start swelling | 4h | [73] |
| 31 | MV_RT | Leukocyte movement rate | 1.44 μm/min | [74, 75] |
| 32 | N_H_KILL | Neutrophil-hyphae killing probability | 22.71% | [76,77,78,79] |

| | | | | |
|---|---|---|---|---|
| 33 | MA_H_KILL | Macrophage-hyphae killing probability | 9.85% | [80,81] |
| 34 | N_PHAG | Probability of neutrophil to phagocytose swollen conidia | 14.72% | [79] |
| 35 | MA_PHAG | Probability of macrophage to phagocytose swollen conidia | 90.55% | [82,83] |
| 36 | MA_MAX_CONIDIA | Maximum number of phagocytosed conidia inside a macrophage | 18 | [83] |
| 37 | N_MAX_CONIDIA | Maximum number of phagocytosed conidia inside a neutrophil | 3 | [83,84] |
| 38 | E_INT | Epithelial cell-Aspergillus (swollen conidia or hyphae) interaction probability | 4.49% | [85] |
| 39 | PHAG_KILL | Leukocyte probability to kill ingested conidia | 1.28% | [86] |
| 40 | MA_HALF_LIFE | Macrophage half-life | 24h | [87] |
| 41 | N_HALF_LIFE | Neutrophil half-life | 6h | [88] |
| 42 | PR_NET | Probability that an activated neutrophil will undergo NETosis instead of apoptosis | 30% | [89] |
| 43 | H_VOL | Hyphae volume | $1.06 \times 10^{-12}$ L | [90,91,92, 93] |
| 44 | SEPATE_L | Length of hyphal segments between septa | 40 μm | [92,93] |
| 45 | CONIDIA_VOL | Swollen conidia volume | $4.84 \times 10^{-14}$ L | [91] |
| 46 | MA_VOL | Macrophage volume | $4.85 \times 10^{-12}$ L | [94] |
| 47 | TURNOVER_RT | Molecule exchange rate between lung and whole-body serum | $0.1823 h^{-1}$ | [95] |
| 48 | MAX_N | Maximum number of neutrophils | 522 | [96, 97, 98, current manuscript] |
| 49 | MIN_N | Minimum number of neutrophils | 15 | |
| 50 | MAX_MA | Maximum number of macrophages | 627 | |
| 51 | MIN_MA | Minimum number of macrophages | 15 | |
| 52 | REC_RT | Leukocyte recruitment rate | $9.77 \times 10^{14}$ | [97] |
| 53 | ATII_NUM | Number of type II pneumocytes | 640 | [99, 100] |
| 54 | AVG_ATI_NUM | Average number of type I pneumocytes | 320 | |
| 55 | AF_INIT_IRON | A. fumigatus initial iron content | $3.83 \times 10^{-18}$ mol | [91,68] |
| 56 | AF_INIT_HEME | A. fumigatus initial heme | $1.03 \times 10^{-18}$ mol | [69] |
| 57 | HEME_UP | Heme uptake rate | $1.56 \times 10^{-3} L \cdot cell^{-1} \cdot h^{-1}$ | [69] |
| 58 | HEME_QTTY | Amount of heme that enters the alveoli upon hemorrhage | $2.07 \times 10^{-16}$ mol | Current manuscript |
| 59 | PR_NET_KILL | Probability that NET will kill a type I epithelial cell | 5.13% | [101] |
| 60 | PR_HYPHAE_KILL | Probability that hyphae will kill a type I epithelial cell | 2.51% | [102] |
| 61 | EPI_INHIB | Type I epithelial cells rate of inhibition of hyphae elongation | 50% | [71] |
| 62 | NET_HALF_LIFE | NET half-life | 3h | [103] |

Notes: Parameters 1-5 were obtained as described [1]. Probabilities of phagocytosis, killing, and interaction refer to the likelihood of an event succeeding in one iteration if the appropriate conditions apply. The maximum number of cells and the average number of epithelial cells is computed over the simulated space. Kcat, Enzymatic turnover number;

Kd, dissociation constant; Km, Michaelis constant; TAFC, triacetylfusarinine C (*Aspergillus* siderophore); Tf, Transferrin.

**Table S2:** Interaction rules in the mathematical model.

| ID | Interaction | Description | Type | Outcome | Reference |
|---|---|---|---|---|---|
| 1 | Macrophage -Neutrophil | Macrophage phagocytose apoptotic neutrophil | Probabilistic: Constant Probability | Phosphatidyl serine receptor activated | [9] |
| 2 | Macrophage -Aspergillus | Macrophage phagocytose swollen conidia | Probabilistic: Constant Probability | Dectin-2 receptor activated; conidia are internalized and subsequently killed | [4,105,106] |
| 3 | Macrophage -Aspergillus | Macrophage kills hyphae | Probabilistic: Constant Probability | Dectin-2 receptor activated; hyphae are killed | [4,80,81] |
| 4 | Macrophage -IL10 | Macrophage with phenotype M1, M2A, M2B, and M2C secrete IL10 | Deterministic: fixed amount | The local concentration of IL10 increases | [6,10,14,21,104, 107] |
| 5 | Macrophage -TGF | Macrophage with phenotype M2C secrete TGF | Deterministic: fixed amount | The local concentration of TGF increases | [6,10,14,21,107] |
| 6 | Macrophage -TNF | Macrophage with phenotype M1 and M2B secrete TNF | Deterministic: fixed amount | The local concentration of TNF increases | [2,3,5,6,10, 14,21,104,107,1 08] |
| 7 | Macrophage -CCL4 | Macrophage with phenotype M1 secretes CCL4 | Deterministic: fixed amount | The local concentration of CCL4 increases | [2,104,107] |
| 8 | Macrophage -CXCL2 | Macrophage with phenotype M1 secretes CXCL2 | Deterministic: fixed amount | The local concentration of CXCL2 increases | [5,104,108] |
| 9 | Macrophage -IL10 | Macrophage primed by IL10 | Probabilistic: constant probability | IL10 Receptor activated | [104,109] |
| 10 | Macrophage -TGF | Macrophage primed by TGF | Probabilistic: constant probability | TGF Receptor activated | [8,9,104] |
| 11 | Macrophage -TNF | Macrophage primed by TNF | Probabilistic: constant probability | TNF Receptor activated | [27,104] |
| 12 | Macrophage -Transferrin | Macrophage import/export iron from/to transferrin | Deterministic: import and export rate are proportional to the external levels of transferrin bound to iron and internal iron levels | Internal and external levels of iron and transferrin-bound iron change. | [64,110,111,112] |
| 13 | Neutrophil-Lactoferrin | Active Neutrophils release lactoferrin | Deterministic: fixed amount | The local concentration of lactoferrin increases | [77,113] |
| 14 | Neutrophil-Aspergillus | Neutrophils phagocytose swollen conidia | Probabilistic: constant probability | Neutrophils become active; swollen conidia are internalized and subsequently killed | [77] |
| 15 | Neutrophil-Aspergillus | Neutrophils kill hyphae | Probabilistic: constant probability | Neutrophils become active; hyphae are killed | [77,105] |
| 16 | Type II Pneumocyte -Aspergillus | Type II Pneumocyte interacts with swollen conidia or hyphae | Probabilistic: initial interaction has a fixed probability. Once the | Type II Pneumocyte becomes cytokine-secreting | [114,115] |

| | | | | | |
|---|---|---|---|---|---|
| | | | interaction is established it is stable | | |
| 17 | Type II Pneumocyte -TNF | cytokine secreting Type II Pneumocyte secrete TNF | Deterministic: fixed amount | The local concentration of TNF increases | [31] |
| 18 | Type II Pneumocyte -CCL4 | Chemokine secreting Type II Pneumocyte secrete CCL4 | Deterministic: fixed amount | The local concentration of CCL4 increases | [116] |
| 19 | Type II Pneumocyte -CXCL2 | Chemokine secreting Type II Pneumocyte secrete CXCL2 | Deterministic: fixed amount | The local concentration of CXCL2 increases | [116] |
| 20 | Type II Pneumocyte -TNF | Type II Pneumocyte is primed by TNF | Probabilistic: constant probability | Type II Pneumocyte becomes cytokine/chemokine secreting | [31] |
| 21 | Aspergillus-TAFC | Aspergillus (hyphae and swollen conidia) with TAFC node "ON" secretes TAFC | Deterministic: fixed amount | The local concentration of TAFC increases | [68,117,118] |
| 22 | Aspergillus-TAFC | Aspergillus with nodes MirB and EstB "ON" import TAFC bound to iron * | Deterministic: proportional to the concentration of TAFC bound to iron | The local concentration of TAFC-bound iron decreases. Internal iron pool increases | [66,117] |
| 23 | TAFC-transferrin | TAFC sequester iron from transferrin bound to iron | Deterministic: Michaelian kinetics | The local concentration of TAFC and Tf-iron decreases Local levels of TAFC-iron and free-transferrin increases. | [65,118] |
| 24 | Lactoferrin-transferrin | Lactoferrin sequester iron from transferrin bound to iron | Deterministic: Michaelian kinetics | The local concentration of lactoferrin and Tf-iron decreases. Local levels of lactoferrin bound to iron and free-transferrin increases. | [32,119] |
| 25 | Macrophage -iron | Necrotic macrophage releases its iron content | Deterministic: the whole iron content of the cell is released | The local concentration of iron increases | NA** |
| 26 | Iron-Aspergillus | Dead hyphae release its iron content | Deterministic: the whole iron content of the cell is released | The local concentration of iron increases | NA** |
| 27 | Iron-Iron-transport-molecule | Lactoferrin, Transferrin, or TAFC chelates the whole iron in content of the voxel | Deterministic: the molecules race to chelate the iron in the voxel. The first to be selected chelates the whole iron content or the maximum of its capacity | The local iron concentration decreases to zero. Iron bound to carriers (TAFC-iron, Lactoferrin-iron, or Tf-iron) increase. | NA |
| 28 | Type I Pneumocytes-Aspergillus | Hyphae kill type I pneumocytes. Pneumocytes that | Probabilistic: constant probability to kill | Type I cell dies or is injured | [102] |

| | | | | | |
|---|---|---|---|---|---|
| | | are not killed are injured. | | | |
| 29 | Type I Pneumocytes-Aspergillus | Live Type I pneumocyte decrease hyphae elongation rate | Deterministic: decrease the elongation rate by a fix percentage | Elongation rate decreases | [71] |
| 30 | Type I pneumocyte-Heme | Dead type I pneumocytes releases extracellular heme in the alveolus | Deterministic | Heme become available to interact with neutrophils and Aspergillus. | This work |
| 31 | Type I-Neutrophil | NET kill type I pneumocyte | Probabilistic: constant probability (default scenario).<br>Deterministic: NET kill injured cells (alternative scenario) | Type I cell become dead | [101] |
| 32 | Heme-Aspergillus | Aspergillus uptake Heme | Deterministic: proportional to the Heme concentration | The local heme concentration decrease. Internal Iron and Heme concentration increases | [69] |
| 33 | Heme-Neutrophil | Neutrophils are primed by Heme | Probabilistic: constant probability | Neutrophils become active | [120] |

* MirB and EstB respectively mediate the uptake and hydrolysis of TAFC-iron complex by *Aspergillus.*
** For simplicity, we assume that upon death, host and fungal cells release their iron in free form.
NET, neutrophil extracellular trap; TAFC, triacetylfusarinine C (*Aspergillus* siderophore); Tf, Transferrin
